# AllSome Sequence Bloom Trees

**DOI:** 10.1101/090464

**Authors:** Chen Sun, Robert S. Harris, Rayan Chikhi, Paul Medvedev

## Abstract

The ubiquity of next generation sequencing has transformed the size and nature of many databases, pushing the boundaries of current indexing and searching methods. One particular example is a database of 2,652 human RNA-seq experiments uploaded to the Sequence Read Archive. Recently, Solomon and Kingsford proposed the Sequence Bloom Tree data structure and demonstrated how it can be used to accurately identify SRA samples that have a transcript of interest potentially expressed. In this paper, we propose an improvement called the AllSome Sequence Bloom Tree. Results show that our new data structure significantly improves performance, reducing the tree construction time by 52.7% and query time by 39 - 85%, with a price of up to 3x memory consumption during queries. Notably, it can query a batch of 198,074 queries in under 8 hours (compared to around two days previously) and a whole set of *k*-mers from a sequencing experiment (about 27 mil *k*-mers) in under 11 minutes.

## 1 Introduction

Data structures for indexing and searching of databases have always been a core contribution of algorithmic bioinformatics to the analysis of biological data and are the building blocks of many popular tools [21]. Traditional databases may include reference genome assemblies, collections of known gene sequences, or reads from a single sequencing experiment. However, the ubiquity of next generation sequencing has transformed the size and nature of many databases. Each sequencing experiment results in a collection of reads (gigabytes in size), typically deposited into a database such as the Sequence Read Archive (SRA) [17]. There are thousands of experiments deposited into the SRA, creating a database of unprecedented size in genomics (4 petabases, as of 2016).The SRA enables public access of the database via meta-data queries on the experiments’ name, type, organism, etc. However, efficiently querying the raw read sequences of the database has remained out of reach for today’s indexing and searching methods, until earlier this year [34].

Given a transcript of interest, an important problem is to identify all publicly available sequenced samples which express it. The SRA contains thousands of human RNA-seq experiments, providing a powerful database to answer this question. One approach is to use tools such as [37, 30, 4] to first identify transcripts present in each of the experiments; however, running these tools on a massive scale is time prohibitive (though cloud-enabled tools like Rail-RNA [29] are making inroads). Moreover, they introduce biases and can easily miss a transcript that is supported by the reads. Another approach is to align the SRA reads to the transcript of interest; however, this approach is infeasible for such large datasets [34].

Recently, Solomon and Kingsford proposed the Sequence Bloom Tree (SBT) data structure and demonstrated how it can accurately identify samples that may have the transcript of interest expressed in the read data [34]. SBT was a breakthrough, allowing to query a set of 214,293 transcripts against a database of 2,652 human RNA-seq experiments in just under 4 days. The SBT is not intended to replace more thorough methods, like alignment, but is intended to be complementary, narrowing down the set of experiments for which a more rigorous investigation is needed.

In this paper, we present the AllSome Sequence Bloom Tree (SBT-ALSO), a time and space improvement on the original SBT (denoted by SBT-SK). It combines three new ideas. The first one is a better construction algorithm based on clustering. The second one is a different representation of the internal nodes of the tree so as to allow earlier pruning and faster exploration of the search space. The final one is building a Bloom filter on the query itself. This allows quick execution of queries that are not just transcripts but are themselves large sequencing experiments.

We evaluate SBT-ALSO on the database of 2,652 human RNA-seq sequencing runs used in [34]. SBT-ALSO reduces tree construction time by 52.7%, when given the Bloom filters of the datasets. It reduces query time by 39 - 85%, with a price of up to 3x memory consumption. Notably, it can query a batch of 198,074 queries in under 8 hours, compared to over two days for SBT-SK. It can also query a whole set of *k*-mers from a sequencing experiment (about 27 mil *k*-mers) in under 11 minutes, compared to more than 23 hours by SBT-SK. Our software is open source and freely available via GitHub^1^.

## 2 Related Work

This work falls into the general category of string pattern matching, where we are asked to locate all occurrences of a short pattern in a large text. In many cases, it is useful to pre-process the text to construct an index that will speed up future queries. The *k*-mer-index, trie, suffix tree, suffix array, BWT, and FM-index are examples of such indices [21]. These form the basis of many read alignment tools such as BWA-mem [18] and Bowtie 2 [16]. While many of these approaches are space and time efficient in their intended setting, they can nevertheless be infeasible on terabyte or petabyte scale data. Other approaches based on word-based indices [28, 39] and compressive genomics [20, 38] do not help for the type of data and queries we consider in this paper.

A Bloom filter (BF) is widely used to improve scalability by determining whether the pattern occurs in the text, without giving its location. It is a space efficient data structure for representing sets that occasionally provides false-positive answers to membership queries [3]. For pattern matching, a BF can be constructed for all the constituent *k*-mers (strings of length of *k*) of the text. Then, if a high percentage of a pattern’s constituent *k*-mers match, the text is a potential match and a full search can be performed. BFs are used in several bioinformatics contexts such as assembly [25, 6, 33, 12], to index and compress whole genome datasets [32], and to compare sequencing experiments against whole genomes [35].

When pattern matching against a database of read collections from sequencing experiments, additional factors need to be considered. First, the reads contain sequencing errors. Second, they only represent short fragments of the underlying DNA and are typically much shorter than the pattern. Third, there are many texts, each of which is its own sequencing experiment. The goal is to identify all texts that match the pattern. A simple way to adapt the BF idea to this case is to simply build a BF for every text and check the pattern separately against every text’s BF. A more sophisticated approach builds a tree to index the collection of BFs [8]. This Bloofi data structure was introduced in the context of distributed data provenance, but it was later adapted to the bioinformatics setting by Solomon and Kingsford in [34].

An orthogonal approach is the Bloom Filter Trie (BFT) [13], which works similarly to a trie on the *k*-mers in all the texts. Each leaf contains a bitvector describing the texts in which that *k*-mer appears, and BFs are cleverly used inside the trie to “jump down” ℓ positions at a time, thus speeding up the trie traversal process. The BFT complexity scales up with the number of *k*-mers in the query, while SBT complexity scales up with the number of datasets. Thus the two approaches suggest orthogonal use cases. In particular, the BFT is very efficient for queries that are single *k*-mers, significantly outperforming the SBT. An approach that uses BFT to query longer patterns like the ones we consider in this paper is promising but is not yet available.

There is also a body of work about storing and indexing assembled genomes [19, 26, 2, 24, 10], which is part of the growing field of pangenomics [7]. However, our work relates to the indexing of unassembled data (i.e. reads) as opposed to complete genomes. In addition to the topics specifically mentioned above, there are other studies related to scaling up indexing methods [23, 9], though the list here is in no way complete.

## 3 Technical Background

### Terminology

Let *x* and *y* be two bitvectors of the same length. The bitwise AND (i.e. intersection) between *x* and *y* is written as *x* ∩ *y*, and the bitwise OR (i.e union) is *x* ∪ *y*. A bitvector can be viewed as a set of positions set to 1, and this notation is consistent with the notion of set union and intersection. The set difference of *x* and *y* is written as *x \ y* and can be defined as *x \ y* = *x* AND (NOT *y*). A *Bloom filter* (BF) is a bitvector of length *b*, together with *p* hash functions, *h_1_*,…, *h_p_*, where *b* and *p* are parameters. Each hash function maps a *k*-mer to an integer between 0 and *b* – 1. The empty set is represented as an array of zeros. To add a *k*-mer *x* to the set, we set the position *h_i_*(*x*) to 1, for all *i*. To check if a *k*-mer *x* is in the set, we check that the position *h_i_*(*x*) is 1, for all *i*. In this paper, we assume that the number of hash functions is 1 (see Section 6 for a discussion). Next, consider a rooted binary tree. The parent of a non-root node *u* is denoted as *parent*(*u*), and the set of all the leaves of the subtree rooted at a node *v* is denoted by *leaves*(*v*). Let *lchild*(*u*) and *rchild*(*u*) refer to the left and right children of a non-leaf node *u*, respectively.

### SBT

Let *Q* be a non-empty set of *k*-mers, and let *B* be a *k*-mer BF. Given 0 ≤ *θ* ≤ 1, we say that *Q θ*-matches *B* if |{*x* ∈ *Q*: *x* exists in *B*}|/|*Q*| ≥ *θ*. That is, the percentage of *k*-mers in *Q* that are also in *B* (including false positive hits) is at least *θ*. Solomon and Kingsford consider the following problem. We are given a database *D* = {*D*_1_,…, *D_n_*}, where each *D_i_* is a BF of size *b*. The query is a *k*-mer set *Q*, and the result of the query should be the set {*i*: *Q θ*-matches *D_i_*}. The goal is to build a data structure that can construct an index on *D* to support multiple future queries.

We make a distinction between the abstract data type that Solomon and Kingsford propose for the problem and their implementation of it. We call the first SBT, and the second SBT-SK (note that in [34] no distinction is made and SBT refers to both). A rooted binary tree is called a *Sequence Bloom Tree* (SBT) of a database *D* if there is a bijection between the leaf nodes and the elements of *D*. Define *B*_∪_(*u*) for a leaf node *u* as its associated database element and *B*_∪_(*u*) for an internal node as 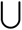*_i∈leaves(*u*)_ B*_∪_(*i*). Note that *B*_∪_(*u*) of an internal node *u* can be equivalently defined as *B*_∪_(*lchild*(*u*)) ∪ *B*_∪_(*rchild*(*u*)). Each node *u* then represents the set of database entries corresponding to the descendant leaves of *u*. Additionally, the SBT provides an interface to construct the tree from a database, to query a *k*-mer set against the database, and to insert/delete a BF into/from the database. An example of an SBT is shown in Figure 2.

### SBT-SK

We call the implementation of the SBT interface provided in [34] as SBT-SK. In SBT-SK, each node *u* is stored as a compressed version of *B*_∪_(*u*). The compression is done using RRR [31] implemented in SDSL [11], which allows to efficiently test whether a bit is set to 1 without decompressing the bitvector. To *insert* a BF *B* into a SBT *T*, SBT-SK does the following. If *T* is empty, it just adds *B* as the root. Otherwise, let *r* be the root. If *r* is a leaf, then add a new root *r’* that is the parent of *B* and *r* and set *B*_∪_(*r’*) = *B*_∪_(*r*)∪*B*. Otherwise, take the child *v* of *r* that has the smallest Hamming distance to *B*, recursively insert *B* in the subtree rooted at *v*, and update *B*_∪_(*r*) to be *B*_∪_(*r*) ∪*B*. Note that because RRR compressed bitvectors do not support bitwise operations, each bitvector must be first decompressed before bitwise operations are performed and then recompressed if any changes are made. The running time of an insertion is proportional to the depth of the SBT. To *construct* the SBT for a database, SBT-SK starts with an empty tree and inserts each element of the database one-by-one. Construction can take time proportional to *nd*, where *d* is the depth of the constructed SBT. The left panel of Figure 1 provides an example of the construction algorithm. To *query* the database for a *k*-mer set *Q*, SBT-SK first checks if *Q θ*-matches the root. If yes, then it recursively queries the children of the root. When the query hits a leaf node, it returns the leaf if *Q θ*-matches it.

**Fig. 1.**
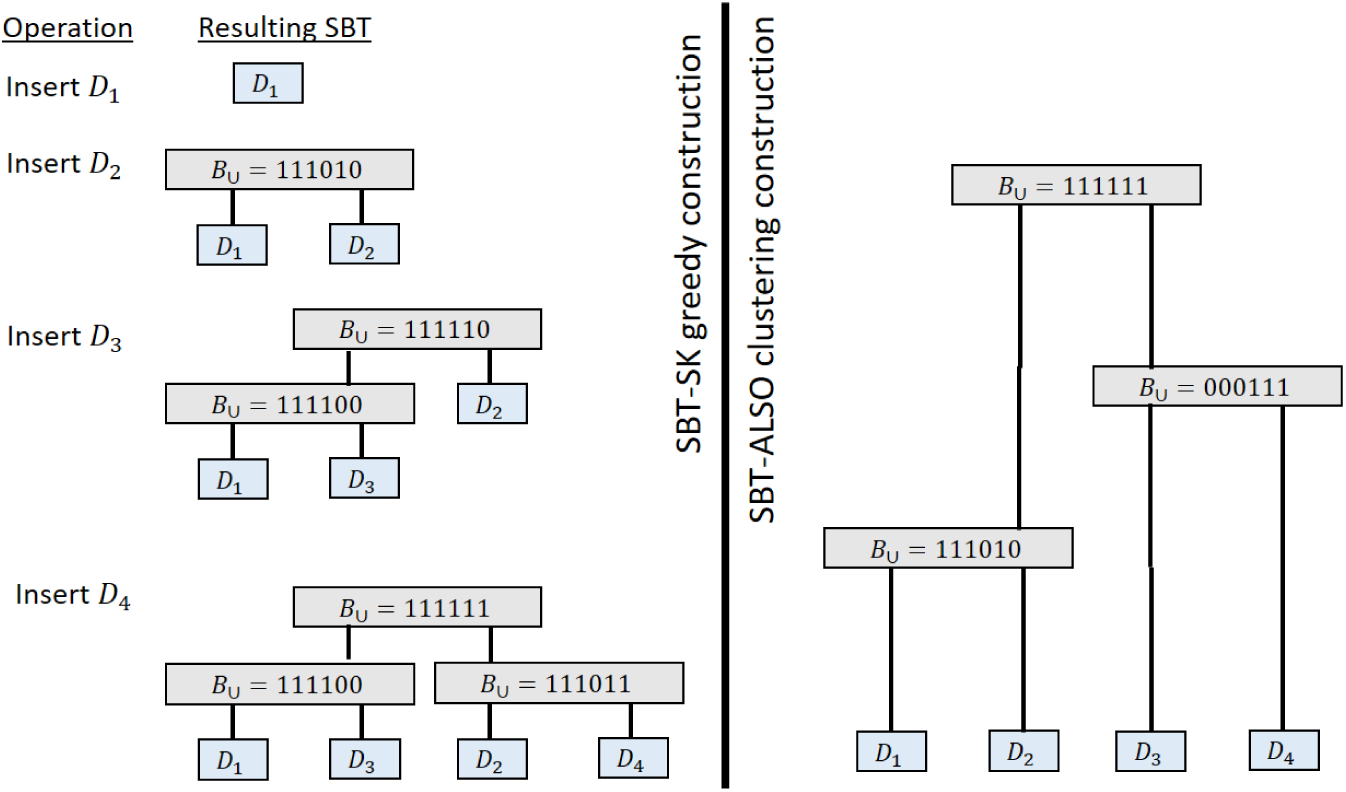
Example of the SBT-SK and SBT-ALSO construction algorithms for the database *D* = {*D*_1_ = 111000, *D*_2_ = 111010, *D*_3_ = 000100, *D*_4_ = 000011}. Leaves shown in blue, internal nodes in gray. In this example, the dataset can be partitioned into two types: 000xxx and 111xxx, based on the first 3 bits. In the SBT-SK construction, after the first two experiments are inserted (both of type 111xxx), they are destined to be in the two different sides of the tree (regardless of future insertions). Any future 111xxx type query will have to examine all the nodes. The SBT-ALSO construction, on the other hand, groups together the experiments so that future 000xxx type or 111xxx type queries will have to examine only about half the nodes of the tree.

Since SBT is designed to work on very large databases, its implementation should avoid loading the database into memory. In SBT-SK, each *B*_∪_(*u*) is stored on disk and only loaded into memory when *u* is being *θ*-matched by a query. When there are multiple queries to be performed, SBT-SK will *batch* them together so that the *θ*-matching of multiple queries to the same node will be performed simultaneously. Hence, each node needs to be loaded into memory only once per batch. We implement the same strategy in SBT-ALSO.

## 4 Methods

We propose the AllSome SBT as an alternative implementation of the SBT abstract data type. In this section, we describe the construction and query algorithms. Insertion and deletion algorithms are the same as in SBT-SK, though some special care is needed. For completeness, they are described in the full version [36].

### 4.1 AllSome node representation and regular query algorithm

Define the intersection of leaves in the subtree rooted at a node *u* as 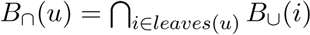 Intuitively, we can partition the 1 bits of *B*_∪_(*u*) into three sets: *B_all_*(*u*), *B_some_*(*u*), and *B*_⋂_(*parent*(*u*)). *B_all_*(*u*) are the bits that appear in all of *leaves*(*u*), excluding those in all of *leaves*(*parent*(*u*)). *B_some_*(*u*) are the bits in some of *leaves*(*u*) but not in all. Both sets therefore exclude bits present in *B*_⋂_(*parent*(*u*)). Formally, define

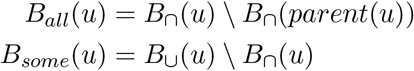

At the root *r*, define *B*_⋂_(*parent*(*r*)) = 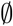. *B_all_*(*u*) and *B_some_*(*u*) are stored using two bitvectors of size *b* compressed with RRR. *B*_∪_(*u*) and *B*_⋂_(*u*) are not explicitly stored. We refer to this representation of the nodes using *B_all_* and *B_some_* as the ALLSOME representation (Figure 2 gives an example).

**Fig. 2.**
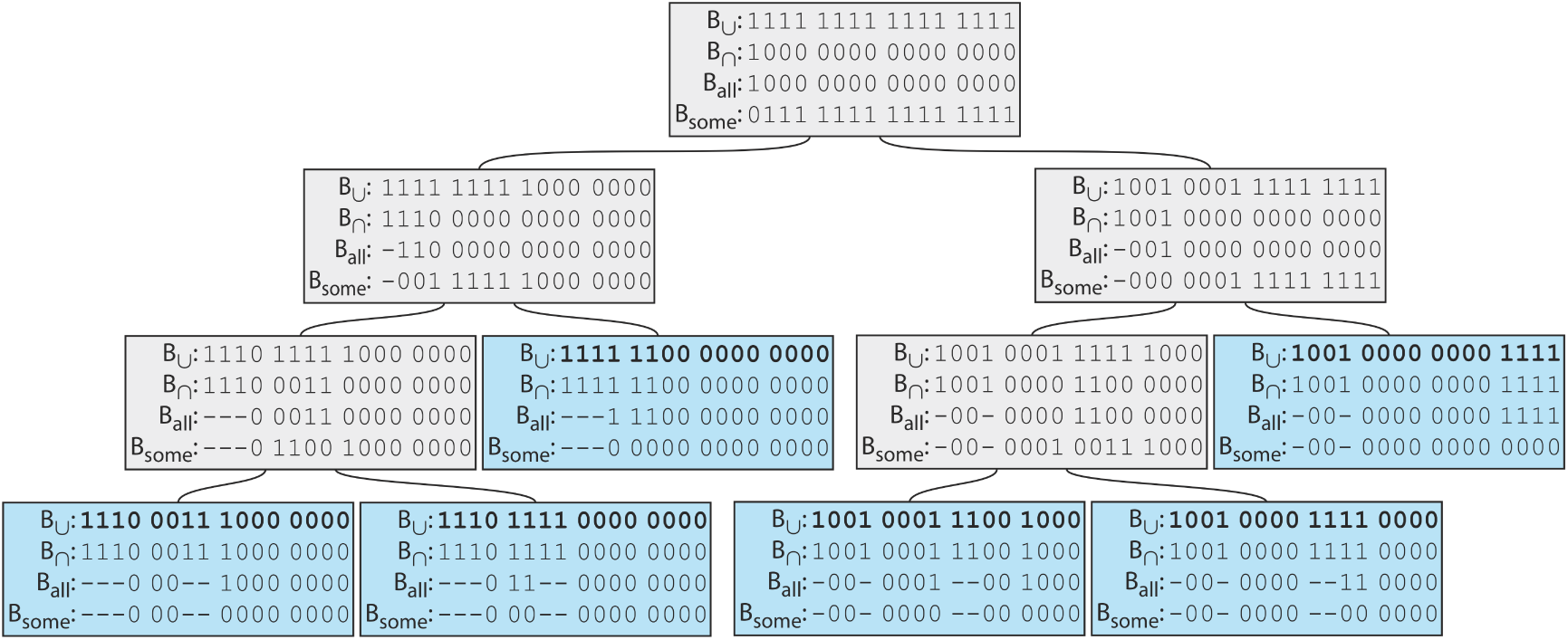
Example SBT on *D* = {1110001110000000, 1110111100000000, 1111110000000000, 1001000111001000, 1001000011110000, 1001000000001111}. Leaves shown in blue, internal nodes in gray. In SBT-ALSO, only *B_all_* and *B*_some_ are explicitly stored, while in SBT-SK, only *B*_∪_ is stored. Bits present in *B_all_* at one node are shown as hyphens(’-’) in the *B_all_* and *B_some_* of its descendants, but in the actual SBT-ALSO data structure they are zeros.

When we receive a query *k*-mer set *Q*, we hash each *k*-mer to determine the list of BF bits corresponding to *Q*. These are a multi-set of position indices (between 0 and *b*–1), stored as an array. We call these the list of *unresolved* bit positions. We also maintain two counters: the number of bit positions that have been determined to be 1 (*present*), and the number of bit positions determined to be 0 (*absent*). These counters are both initially 0. The query comparison then proceeds in a recursive manner. When comparing *Q* against a node *u*, each unresolved bit position that is 1 in *B_all_*(*u*) is removed from the unresolved list and the present counter is incremented. Each unresolved bit position that is 0 in B*_some_*(*u*) is removed from the unresolved list and the absent counter is incremented. If the present counter is at least *θ*|Q|, we add leaves(*u*) to the list of θ-matches and terminate the search of *u*’s subtree. If the absent counter exceeds (1 – θ) |*Q*|, we realize that *Q* will not *θ*-match any of the leaves in the subtree rooted at *u* and terminate the search of *u*’s subtree. If neither of these holds, we recursively pass the two counters and the list of unresolved bits down to its children. When we reach a leaf, the unresolved list will become empty because *B_some_* is empty at a leaf, and the algorithm will necessarily terminate.

The idea behind the ALLSOME representation is that in a database of biologically associated samples, there are many *k*-mers that are shared between many datasets. In the SBT-SK representation, a query must continue checking for the presence of these *k*-mers at every node that it encounters. By storing at *u* all the bits that are present in all the leaves of its subtree, we can count those bits as resolved much earlier in the query process – limiting the amount of bit look-ups performed. Moreover, we will often prune the search space earlier and decrease the number of bitvectors that need to be loaded from disk. A query that matches all the leaves of a subtree can often be resolved after just examining the root of that subtree. In the extreme case, the number of nodes examined in a search may be less than the number of database entries that are matched.

A second important point is that the size of the uncompressed bitvectors at each node is now twice as large as before. Because query time has a large I/O component, this has potential negative effects. Fortunately, we observe that the compressed size of these bitvectors is roughly proportional to the number of 1s that are contained. By defining the ALLSOME reprsentation as we do, the number of 1s in total in *B_all_*(*u*) and *B_some_*(*u*) is no more than the number of ones in *B*_∪_(*u*). Moreover, because we exclude *B*_⋂_(*u*) from all of *u*’s descendants, the number of 1s is less.

### 4.2 Construction algorithm

Except for large queries or large batches of queries, the running time of the query algorithm is dominated by the I/O of loading bitvectors into memory [34]. If the number of leafs that the query *θ*-matches is localized within the same part of the SBT, then fewer internal nodes have to be explored and, hence, fewer bitvectors have to be loaded into memory. The SBT-SK construction algorithm is greedy and sensitive to the order in which the entries are inserted into the tree, which can lead to trees with poor localization (see example in Figure 1).

To improve the localization property of the tree, we propose a non-greedy construction method based on agglomerative hierarchical clustering [14]. Every *D_i_* is initially its own SBT, with its *B*_∪_ loaded into memory. At every step, two SBTs are chosen and joined together to form a new SBT. The new SBT has a root node *r* with the left and right subtrees corresponding to the two SBTs being joined. *B*_∪_(*r*) is computed as *B*_∪_(*lchild*(*r*)) ∪ *B*_∪_(*rchild*(*r*)). To choose the pair of SBTs to be joined, we choose the two SBTs that have the smallest Hamming distance between the *B*_∪_ of their roots. The right panel of Figure 1 shows how our construction algorithm works.

Since each *B*_∪_ is a large bitvector, computing and maintaining the pairwise distances between all pairs is computationally expensive. Instead, we use the following heuristic. We fix a number *b*′≪ *b* (e.g. *b*′ = 10^5^≪10^9^ = *b*) and then extract *b*′ bits from each *D_i_*, starting from a fixed but arbitrary offset. We then run the above clustering algorithm on this smaller database of extracted bitvectors.

The resulting topology is then extracted and used for constructing the *B_all_* and *B_some_* bitvectors for all the nodes. We process the nodes in a bottom-up fashion. Initially, for all leaves *u*, we set *B_all_*(*u*) = *B*_∪_(*u*) and *B_some_*(*u*) = 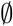. For the general case, consider an internal node *u* whose children ℓ and *r* have already been processed. All bits that are set in both *B_all_*(*l*) and *B_all_*(*r*) go into *B_all_*(*u*):

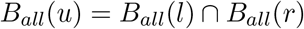

Additionally, the *B_all_* bits of ℓ and *r* must exclude those that are set in the parent *B_all_*(*u*). After computing *B_all_*(*u*), we can unset these bits:

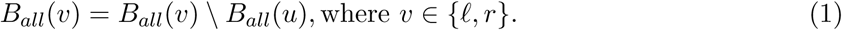

Note that this is the only necessary update to the bitvectors of nodes in the subtree rooted in ℓ or *r*. Next, we must compute *B_some_*(*u*), which is the set of bits that exist in some of *u*’s children nodes but not all:

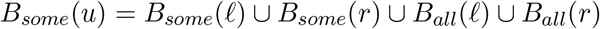
Note that here we are using the *B_all_* after the application of Equation (1). This completes the necessary updates to the tree for a node *u*. These updates can be efficiently computed using bitwise operations on uncompressed bitvectors, so we keep them uncompressed in memory and only compress them when they are written to disk and are no longer needed. The total time for construction is proportional to *n* and not to *nd*, as with SBT-SK. For completeness, we provide a more formal algebraic derivation of the update formulas in the full version [36].

### 4.3 Large query algorithm

The “regular query” algorithm (Section 4.1) is designed with relatively small queries in mind (e.g. thousands of *k*-mers from a transcript). However, after performing a new sequencing experiment, it might be desirable to query the database for other similar samples. In such cases, the query would itself be a whole sequencing experiment, containing millions of *k*-mers. Our experimental results show that neither SBT-SK nor our own regular query algorithm is efficient for these large queries.

While for small queries, the running time is dominated by the I/O of loading bitvectors into memory, for large queries, the time taken to look up the query *k*-mers in the *B_all_* and *B_some_* of a node becomes the bottleneck. Let *B_Q_* be the Bloom filter of size *b* for the *k*-mers in the query *Q*. We propose an alternate “large query” algorithm that can be used whenever the number of *k*-mers in the query exceeds some pre-defined user threshold. This large query algorithm is identical to the regular one except in the way that the unresolved list is maintained and updated. The basic idea is that instead of checking each *k*-mer in *Q* one-by-one, we can do bitwise comparisons using *B_Q_*. Assume for the moment that there are no two *k*-mers in *Q* that hash to the same position (recall that our BFs have only one hash function). In this case, the list of unresolved bit positions can be represented as the set of 1 positions in *B_Q_*. At a node *u*, we first increment the *present* counter by the number of ones in *B_Q_*⋂*B_all_*(*u*) and update the unresolved bit positions to be 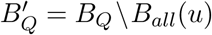. Then we increment the *absent* counter by the number of ones in 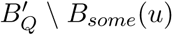 and update the unresolved bit positions to be 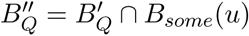. If the counters do not exceed their respective thresholds, then we pass them and the remaining unresolved bits 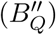 down to the children.

When there are *k*-mers that hash to the same bit positions, the above algorithm can still be used as a heuristic. In fact, it can be shown that the hits returned by the above heuristic algorithm are always a subset of the hits that are returned by an exact algorithm, since the heuristic’s counter values are never greater than those of the exact algorithm. But, we can obtain an exact algorithm by modifying the above heuristic to also maintain a list of bit positions that have multiple *k*-mers hashing to them. An entry of the list is a bit position and the number of *k*-mers that hash to it. Whenever we make a bitwise comparison involving *B_Q_*, this list is scanned to convert numbers of bits to numbers of *k*-mers. When the list is small, this exact algorithm should not be significantly slower than the heuristic one.

Unfortunately, computing bitwise operations cannot be efficiently done on RRR compressed bitvectors. To support the large query algorithm, the bitvectors are compressed using the Roaring [5] scheme (abbreviated ROAR). Roaring bitmaps are compressed using a hybrid technique that allows them to efficiently support set operations on bitvectors (intersection, union, difference, etc). However, we found that they generally do not compress as well as RRR on our data, leading to longer I/O times. In cases where both small and large queries are common, and query time is more important than disk space, both a ROAR and an RRR compressed tree can be maintained.

## 5 Results

We implemented SBT-ALSO, building on the SBT-SK code base [1]. Solomon and Kingsford already explored the advantages, disadvantages and accuracy of the SBT approach as a way of finding experiments where the queried transcripts are expressed [34]. Since SBT-ALSO gives identical query results as SBT-SK, we therefore focus our evaluation on its resource utilization. We used the same dataset for evaluation as in [34]. This is the set of 2,652 runs representing the entirety (at the time of [34]) of human RNA-seq runs from blood, brain, and breast tissues at the SRA, excluding those sequenced with SOLID. In [34], each sequencing run was converted to a *k*-mer Bloom filter (*b* = 2·10^9^, *k* = 20) by the Jellyfish *k*-mer-counting software (containing *k*-mers that occur greater than a file-dependent threshold, typically at least 3 occurrences). We downloaded these BFs from [1] and used them as our database. Per the results of [34], this Bloom filter size leads to a false positive rate of 0.5 for an individual Bloom filter. We performed experiments on an OpenStack instance with 12 vCPUs (Intel Xeon E312xx), 128 GB memory, and 4 TB network-mounted disk storage.

To choose the appropriate number of bits to use for clustering (*b*′), we randomly sampled 5,000 bitvector pairs from the dataset and computed their pairwise distances. We then computed distances for the same pairs using only *b*′ bits, for various values of *b*′. The two distance metrics showed a high correlation (*r*^2^ = 0.9999874) for *b*′ = 500, 000.

We then constructed SBT-SK and SBT-ALSO, as well as two other trees to help us separate out the contributions of the clustering algorithm from the ALLSOME representation. These two trees are SBT-SK+CLUST, which uses the *B*_∪_ node representation of SBT-SK but the SBT-ALSO clustering construction, and SBT-SK+AS, which uses the greedy construction of SBT-SK but the ALLSOME node representation of SBT-SK.

First, we compared the space and time used to construct SBT-SK and SBT-ALSO (Table 1). SBT-ALSO reduces the tree construction time by 52.7% and resulting disk space by 11.4%. It requires twice as much intermediate space, due to maintaining two uncompressed bitvectors for each node instead of just one.

**Table 1.** Construction time and space. Times shown are wall-clock times. A single thread was used. Note the SBT-SK tree that was constructed for the purposes of this Table differs from the tree used in [34] and in our other experiments because the insertion order during construction was not the same as in [34] (because it was not described there).

|  | SBT-SK | SBT-ALSO |
| --- | --- | --- |
| construction of tree topology (i.e. clustering) | N/A | 27m |
| construction of internal nodes | 56h 54m | 26h 3m |
| peak memory usage | 726 MB | 908 MB |
| temporary disk space | 1,235 GB | 2,469 GB |
| final disk space | 200 GB | 177 GB |

To study the regular query performance, we downloaded all known transcripts at least *k* bases long (198,074 of them) from Gencode (ver. 25). We then queried several subsets of transcripts against both trees, and measured the number of nodes examined for each query (Figure 3) as well as the running time (Table 2). The results of all query experiments in this paper were verified to be equivalent between the tested data structures. SBT-ALSO reduces the runtime by 39 - 85%, depending on the size of the batch, likely due to the fact that the number of nodes examined per query is reduced by 52.7%, on average. Notably, SBT-ALSO was able to query a very large batch (198,074 queries) in under 8 hours, while SBT-SK took over 2 days. SBT-ALSO uses more memory than SBT-SK on larger batches.

**Fig. 3.**
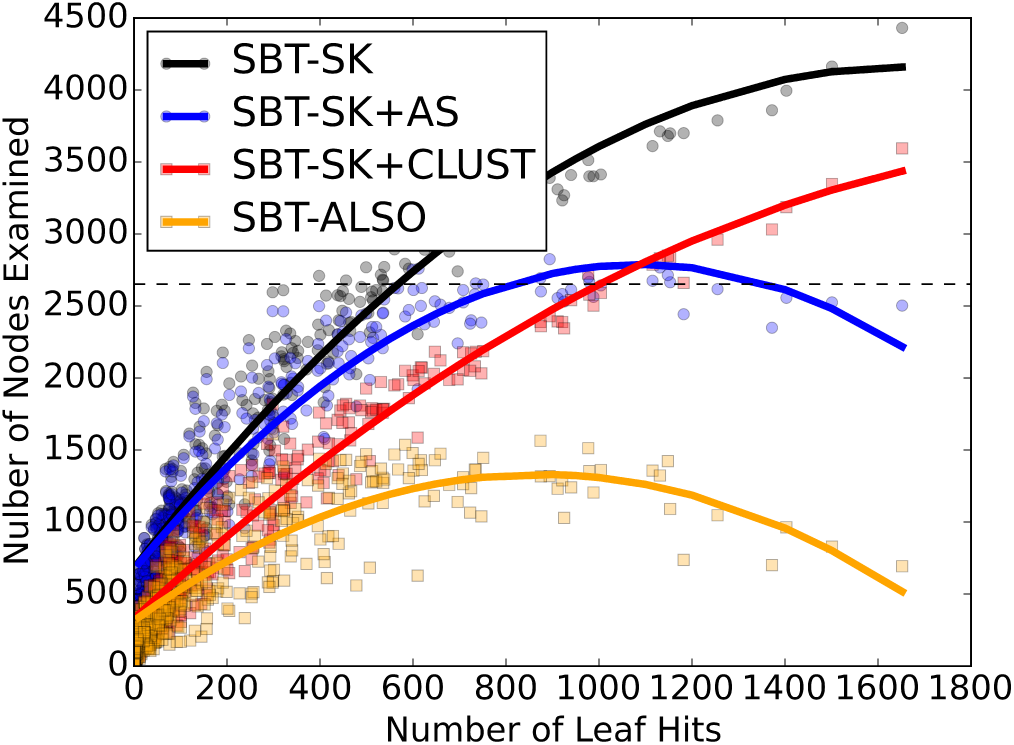
Number of nodes examined per query for SBT-SK, SBT-ALSO, as well two intermediate SBTs. A set of 1,000 transcripts were chosen at random from Gencode set, and each one queried against the four different trees. A dot represents a query and shows the number of matches in the database (x-axis) compared to the number of nodes that had to be loaded from disk and examined during the search (y-axis). For each tree (color), we interpolated a curve to show the pattern. The dashed horizontal line represents the hypothetical algorithm of simply checking if the query *θ*-matches against each of the database entries, one-by-one. For *θ*, we used the default value in the SBT software (*θ* = 0.9).

**Table 2.** Query wall-clock run times and maximum memory usage, for batches of different sizes. For the batch of 1,000 queries, we used the same 1,000 queries as in Figure 3. For the batch of 100 queries, we generated three replicate sets, where each set contains 100 randomly sampled transcripts without replacement from the 1,000 queries set. For the batch of 10 queries, we generated 10 replicate sets by partitioning one of the 100 query sets into 10 sets of 10 queries. For the batch of 1 query, we generated 50 replicate sets by sampling 50 random queries from Gencode set. The shown running times are the averages of these replicates. For *θ*, we used the default value in the SBT software (*θ* = 0.9).

| # query | SBT-SK | SBT-SK+AS | SBT-SK+CLUST | SBT-ALSO |
| --- | --- | --- | --- | --- |
| 1 | 1.2m / 301MB | 2.7m / 301MB | 0.9m / 299MB | 0.5m / 301MB |
| 10 | 4m / 305MB | 8.3m / 319MB | 3.3m / 304MB | 2m / 313MB |
| 100 | 7.7m / 315MB | 13.7m / 346MB | 6.5m / 317MB | 4.7m / 353MB |
| 1,000 | 25.5m / 420MB | 20.8m / 575MB | 17.3m / 418MB | 8.3m / 639MB |
| 198,074 | 3082m / 22GB | 1286m / 51GB | 1910m / 23GB | 463m / 63GB |

To study the performance of the large query algorithm, we selected an arbitrary run from our database (SRR806782) and used Jellyfish [22] to extract all 20-mers that appear at least three times. These 27,546,676 *k*-mers formed one query. In heuristic mode, the large query algorithm was 22 times faster than the regular one, but only detected 47 hits, which is a subset of the 50 hits by regular algorithm (Table 3). In the exact mode, the large query algorithm recovered all the hits (as expected) and was 18 times faster. Compared to SBT-SK, it was 155 times faster.

**Table 3.** Performance of different trees and query algorithms on a large query. We show the performance of SBT-SK and three query algorithms using SBT-ALSO compressed with ROAR: the regular algorithm, the large exact algorithm, and the large heuristic algorithm. We show the wall-clock run time and maximum RAM usage. We used *θ* = 0.8 for this experiment. The ROAR compressed tree was 190 GB (7.3% larger than the RRR tree).

|  | SBT-SK | SBT-ALSO |  |  |
| --- | --- | --- | --- | --- |
|  | regular alg | regular alg | large exact alg | large heuristic alg |
| query time | 1397m 18s | 195m 33s | 10m 35s | 8m 32s |
| query memory | 2.3 GB | 4.7 GB | 1.3 GB | 1.2 GB |

The clustering construction, even without the AllSome representation, significantly reduces the number of nodes that need to be examined per query (36.5% on average when comparing SBT-SK to SBT-SK+CLUST in Figure 3). The improvement seems to be uniform regardless of the number of leaf hits. As expected, this leads a significant improvement to the running time (19-32%, Section 5).

The ALLSOME representation, without the clustering construction, also gives the benefit of allowing earlier query resolution, but the effect only becomes pronounced for queries that hit a lot of leaves. For instance, queries that hit more than 800 leaves examined 27.4% less nodes in SBT-SK+AS then in SBT-SK. In the extreme case, there are seven queries out of 1,000 where the number of nodes examined is less than the number of leaf hits, something that is not possible with SBT-SK. However, the benefits of clustering construction and ALLSOME representation are synergistic: the multiplicative effect of their individual contributions (42.3% decrease in number of examined nodes) is less than the observed effect of their combined contributions (52.7%). In terms of the running time performance, the ALLSOME representation incurs the overhead of making two queries per active bit, instead of just one. This is more than compensated by a decrease in the amount of active bits when the tree is clustered well. But, as the SBT-SK+AS column of section 5 shows, the running time can actually deteriorate when the tree is not clustered.

## 6 Discussion

In this paper we present an alternate implementation of the SBT that provides substantial improvements in query and construction time. We are especially effective for large batches of queries (6 times faster) or for large queries (155 times faster). Solomon and Kingsford make a convincing case that an efficient SBT implementation translates to an efficient and accurate solution to the broader problem of identifying RNA-seq samples that express a transcript of interest. They study the best parameter values of SBT (*θ, k, b, p*) to achieve accuracy and speed for the broader problem. The focus of this paper is on improving resource performance, and hence we do not revisit these questions, however, a more thorough exploration of the biological questions that the SBT can answer will be important moving forward.

The implications of using the SBT for queries which are themselves sequencing experiments were not explored in SBT-SK or here. The BFT [13], if adapted to multi-*k*-mer queries with *θ*-matching, could prove to be powerful in this context. In general, the question of whether the percentage of matching *k*-mers is a good metric for comparing sequencing experiments is still open, and more investigation into how to best measure similarity is needed (e.g. see [27]). However, our large query algorithm opens the door for efficiently exploring the parameter space of *k*-mer-based approaches.

In contrast to SBT-SK, we do not currently support multiple hash functions. For the type of application considered in this paper, [34] demonstrated that one hash function is optimal. Yet, there may be other applications where multiple hash functions offer advantages. This may make SBT-ALSO, in its current state, less broadly applicable then SBT-SK. However, multiple hash functions could be implemented within the AllSome representation using partitioned Bloom filters (where each hash function maps to a different bit array) [15]. This remains as future work.

## Footnotes

1 SBT-ALSO GitHub repository: https://github.com/medvedevgroup/bloomtree-allsome

## Acknowledgements

This work has been supported in part by NSF awards DBI-1356529, CCF- 551439057, IIS-1453527, and IIS-1421908 to PM.

## A Appendix

### A.1 Insertion

If a tree is modified by the addition (or removal) of a leaf, the only nodes for which *B*_∪_ and *B*_∩_ can change are along the path from the leaf to the root. This fact, along with the definitions of *B_all_* and *B_some_*, shows that it is sufficient to only consider changes in *B_some_* along that path, and in *B_all_* along that path and the siblings of those nodes.

To insert a new Bloom filter *B*, we follow the same strategy as SBT-SK. We insert *B* starting at the root and recursively pass it down to the child u that has the smallest Hamming distance between *B*_∪_(*u*) and *B*. Though *B*_∪_(*u*) is not explicitly stored in the SBT-ALSO, it can be recovered on the fly using the equations:

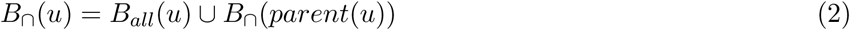

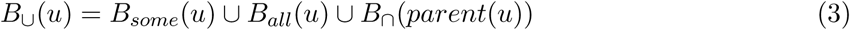
As we proceed down the tree, we must also update the appropriate bitvectors. Consider the insertion of *B* into the subtree rooted at a node *u*. We inductively assume that the bitvectors of nodes outside the subtree rooted at *u* have already been updated, that the bitvectors of nodes inside this subtree have been unchanged, and that *B*_∩_(*parent*(*u*)) is available in memory. We use the superscript *new* to denote the bitvectors of the nodes after *B* is recursively passed down to one of the child’s subtrees.

At *u*, observe that 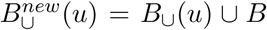 and 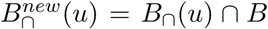. This formula, together with *B*_∩_(*parent*(*u*)), is used to update *B_all_*(*u*) and *B_some_*(*u*), using their corresponding definitions. Assuming without loss of generality that *B* will be passed down to the left child of *u*, the only other node that needs to be updated is the right child. Even though *B*_∪_(*rchild*(*u*)) and *B*_∩_(*rchild*(*u*)) remain unchanged, we need to update 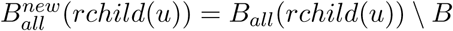.

### A.2 Deletion

Consider the deletion of an entry from the database. Let *v* be the leaf representing the deleted entry, and let *v*′ be its sibling. We set *parent*(*v*′) = *parent*(*parent*(*v*′)) and delete *v* and *parent*(*v*) from the tree, Next, we need to update the bitvectors of the tree.

Let *p* be the path from the root down to *v*′. Let *p*′ be the nodes of *p* along with the children of nodes in *p*. We use the superscript *new* to denote the bitvectors after the deletion, and omit the superscript to indicate bitvectors prior to the deletion. We will update the bitvectors in three passes. In the first pass, we will go down from the root to recover *B*_∩_ and *B*_∪_ for nodes in *p*′ and store them in active memory. In the second pass, we will go up from *v*′ and use the output of the first pass to calculate 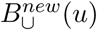 and 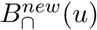 for nodes in *p*. Note that 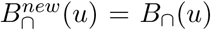 and 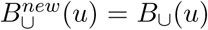 for nodes not in *p*. In the the third pass, will go up from *v'* and use the output of the second pass to calculate 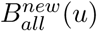 and 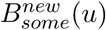 for all nodes on *p*′.

In the first pass, we can recover *B*_∩_ using the equations Equations (2) and (3) above. In the second pass, we can compute (going up from the leaf)

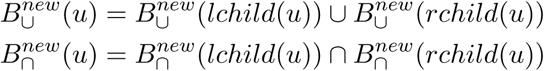
In the third pass, we can compute:

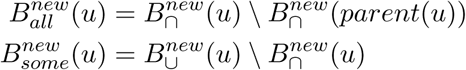
We note that with a smart implementation, the second and third pass can be combined and the computation of *B*_∪_(*u*) in the first pass can be done instead on the fly in the second pass. The above algorithm also requires maintaining *O*(*d*) bitvectors in memory, where *d* is the depth of the tree. If memory is limited, then it is possible to read and write the bitvectors to disk for each node as it is being covered in a pass.

The running time of both an insertion or deletion is on the order of the depth of the tree. Performing an insertion/deletion requires performing bitwise operation on bitvectors, which can be done efficiently on a ROAR compressed tree or an uncompressed tree. If RRR is being used, then, similar to SBT-SK, we need to uncompress nodes before processing them and recompress them after.

Finally, we note that if there are many modifications to the tree, the advantages of the initial clustering construction may dissipate. In this case, the tree can be reconstructed from scratch, incurring a time penalty but reducing the run time of future queries.

### A.3 Formal derivation of update formulas for construction

In Section 4.2, we presented the update rules for constructing *B_all_* and *B_some_* for the internal nodes of a SBT. Here, we give a formal derivation of the rules’ correctness. First, let *DB*(*u*) denote the database entries corresponding to the descendant leaves of a node *u*. Note that the subtree rooted at *u* is, by definition, an SBT of DB(*u*). At any point of the construction, we will have the invariant that if node *v* was processed then the subtree rooted at *v* is a correct SBT-ALSO for DB(*v*). For the base case, for all leaves *u* we set 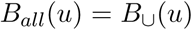. and 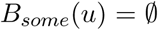. For the general case, consider an internal node *u* whose children have already been processed. We will use the superscript *new* to denote the values of the bitvectors for the new subtree rooted at *u*, to distinguish it from those values passed up inductively from the trees of the children. An important point is that, for a child *v*, 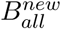 (*v*) may be different from *B_all_*(*v*). This is because once the SBT-ALSO of DB(*v*) is incorporated into the SBT-ALSO of *DB*(*u*), any bits that are set in 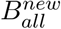. (*u*) need to be unset in 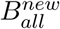. (*v*). Also, observe that for a root r of an SBT-ALSO tree (e.g. *v* in DB(*v*) or *u* in DB(*u*)), *B_all_*(*r*) = *B*_∩_(*r*) and *B*_∪_(*r*) = *B_all_*(*r*) ∪ *B_some_*(*r*). Applying our observations and definition, we can derive formula for 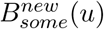, 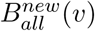, and 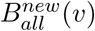.:

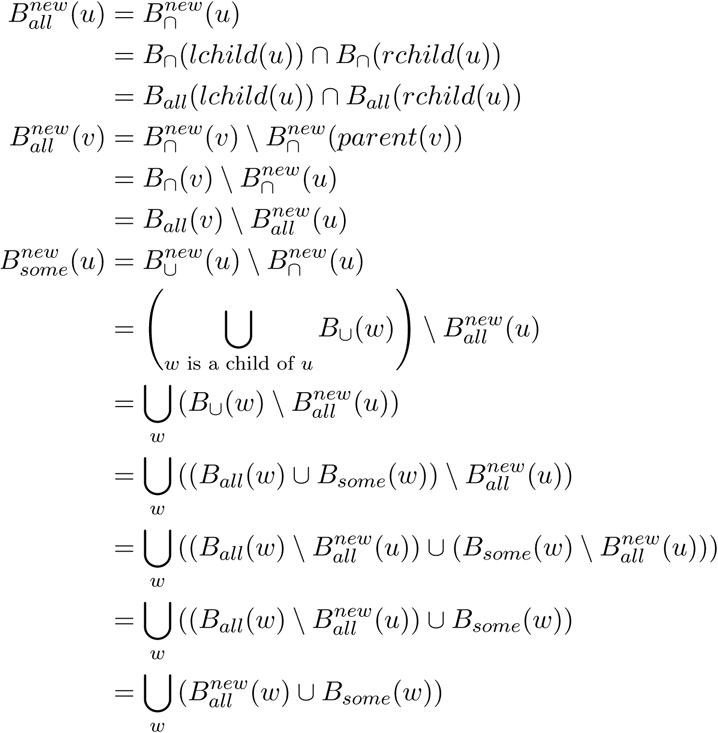

